# Mitochondrial ncRNA LDL-805 declines in alveolar epithelial type 2 cells of chronic obstructive pulmonary disease patients

**DOI:** 10.1101/2024.01.14.575591

**Authors:** Theodore L. Mathuram, Yafei Su, Jonathan E. Bard, Noa A. Perry, Chien Wen Chen, Marisa T. Warren, Phillip A. Linden, Yaron Perry, Maria Hatzoglou, Yun Wu, Anna Blumental-Perry

## Abstract

**Rationale:** We showed that levels of a murine mitochondrial noncoding RNA, *mito-ncR-LDL805*, increase in alveolar epithelial type 2 cells exposed to extracts from cigarette smoke. The transcripts translocate to the nucleus, upregulating nucleus-encoded mitochondrial genes and mitochondrial bioenergetics. This response is lost after chronic exposure to smoke in a mouse model of chronic obstructive pulmonary disease.

**Objectives:** To determine if *mito-ncR-LDL805* plays a role in human disease, this study aimed to (i) identify the human homologue, (ii) test if the smoke-induced response occurs in human cells, (ii) determine causality between the subcellular localization of the transcript and increased mitochondrial bioenergetics, and (iii) analyze *mito-ncR-LDL805* transcript levels in samples from patients with chronic obstructive pulmonary disease.

**Methods:** Levels and subcellular localization of the human homologue identified from an RNA transcript library were assessed in human alveolar epithelial type 2 cells exposed to smoke extract. Lipid nanoparticles were used for nucleus-targeted delivery of *mito-ncR-LDL805* transcripts. Analyses included *in situ* hybridization, quantitative PCR, cell growth, and Seahorse mitochondrial bioenergetics assays.

**Measurements and Main Results:** The levels of human homologue transiently increased and the transcripts translocated to the nuclei in human cells exposed to smoke extract. Targeted nuclear delivery of transcripts increased mitochondrial bioenergetics. Alveolar cells from humans with chronic obstructive pulmonary disease had reduced levels of the *mito-ncR-LDL805*.

**Conclusions:** *mito-ncR-LDL805* mediates mitochondrial bioenergetics in murine and human alveolar epithelial type 2 cells in response to cigarette smoke exposure, but this response is likely lost in diseases associated with chronic smoking, such as chronic obstructive pulmonary disease, due to its diminished levels.

**Impact:** This study describes a novel mechanism by which epithelial cells in the lungs adapt to the mitochondrial stress triggered by exposure to cigarette smoke. We show that a noncoding RNA in mitochondria is upregulated and translocated to the nuclei of alveolar epithelial type 2 cells to trigger expression of genes that restore mitochondrial bioenergetics. Mitochondria function and levels of the noncoding RNA decrease under conditions that lead to chronic obstructive pulmonary disease, suggesting that the mitochondrial noncoding RNA can serve as potential therapeutic target to restore function to halt disease progression.

## Introduction

Exposure to cigarette smoke is a major risk factor for chronic obstructive pulmonary disease (**COPD**), which kills >400,000 Americans each year (1). Lung cells in these individuals lose their ability to repair smoke-induced damage, which results in sustained inflammation, cell death, and emphysema (2,3). The cells that are most affected are type 1 alveolar epithelial cells (**AEC1s**) (4). These cells are replenished by the proliferation and differentiation of type 2 cells (**AEC2s**) to maintain lung homeostasis (5–7). AEC2s are more resistant to damage from cigarette smoke (7–9), but the mechanism is not known. Elucidation of this mechanism could lead to effective therapeutics for COPD.

COPD is associated with severe mitochondrial damage and dysfunction in lung cells (10,11). Although mitochondrial fragmentation in response to cigarette smoke exposure triggers mitophagy, glycolysis, and inflammation in many cell types (12–16), AEC2s respond with increased Krebs cycle activity and elongation of mitochondria to avoid mitophagy (17–19). We began to explore this adaptive mechanism in a previous study and found that a single exposure to cigarette smoke induced an increase in a mitochondrial DNA (**mtDNA**)-encoded noncoding (**nc**) RNA from the light chain D-loop (**LDL**) of microRNA 805 in mouse (*Mus musculus* [*mmu*]) cells, which we call *mmu-mito-ncR-LDL805* (19). This transcript was translocated to the nucleus, coinciding with increased expression of nucleus-encoded genes for mitochondrial proteins (**NeMito**) and mitochondrial bioenergetics. These adaptive responses were lost after chronic exposure to cigarette smoke (19), and we propose that this loss contributes to the development of conditions such as COPD.

*mmu-mito-ncR-LDL805*, to our knowledge, is the only known mtDNA-encoded ncRNA that relocates to the nucleus during stress. This transcript, along with three other mtDNA-encoded ncRNAs, is tightly regulated depending on the energy demand, the ability to supply mitochondrial ribosomes, and whether mtDNA replication-transcription are balanced (20,21). Although *mmu-mito-ncR-LDL805* is a mouse-specific transcript, it contains a 20-nucleotide “functional bit” that is highly conserved among mammals (22). Therefore, a homologous transcript is likely expressed by human cells that may be relevant to the development and progression of smoke-related lung diseases such as COPD. The aim of this study was to identify the human homologue and assess its expression and localization in human AEC2s in response to exposure to cigarette smoke extract. The findings of this study may reveal a novel therapeutic target to protect lung cells and delay disease progression in individuals with COPD.

## Methods

### Cells

MLE12, BEAS-2B, and A549 cells were purchased from ATCC, and primary human AEC2s were from AcceGen. All cells were grown in a humidified incubator supplemented with 5% CO_2_ according to provider instructions.

### RNA Isolation

Was done using an miRNeasy Mini kit (Qiagen) per the manufacturer’s instructions.

### Homologue Identification

To identify the *Homo sapiens* (*hsa*) homologue total RNA was isolated and prepared for illumina sequencing using the NEB Small RNA Library preparation kit. Libraries were sequenced using the illumina NextSeq500 using paried-end 75 cycle sequencing. The FASTQ files were demultiplexed using illumina bcl2fastq, and mapped to the Grch38 reference genome using bowtie2 very-sensititive-local alignments. Alignment files were visualized using the IGV browser to identify the start, stop, strand, and sequence of the target.

### Inhibition of Mitochondrial Transcription by Rhodamine 6G

A549 and BEAS-2B cells were grown in their respective growth media supplemented with 200 μM uridine, 3 μM rhodamine 6G, and 1 mM sodium pyruvate for 72 h, as described before (**23**).

### Mitochondrial Enrichment

Post nuclear fractions were obtained and treated with RNase (to eliminate all but mitochondrial RNAs) as previously described (22).

### RT-qPCR

A high-capacity cDNA reverse transcription kit was used to generate cDNA (Applied Biosystems). qPCRs were performed using TaqMan universal master mix II, no UNG (Life Technologies) and assays LDL1 (RT, TM: CS39QW4; PN4398987), RNU48 (RT, TM: 001006; PN4427975), Oxa1L (hs00192329_m1, Mm01320047_g1), Ndufs8 (hs00159597_m1), Ndufa10 (hs01071117_m1, Mm01290377_g1), Ndufb7 (Mm00788005_s1). Results were calculated using the 2^−ΔΔ*CT*^ method.

### FISH Combined with Immunofluorescence

Detection of *hsa-mito-ncR-LDL805* by fluorescence *in situ* hybridization (FISH) was performed using the miRNAScope HD detection reagent assay (ADC), probe 5’-CGACATCTGGTTCCTACTTC-3’ and combined with immunofluorescence for citrate synthase according to ADC protocols, except that AMP 5 step was shorten to 15 min.

### Cigarette Smoke Extract

Cigarette smoke extract was prepared using research-grade 1R5F cigarettes (Kentucky Tobacco and Health Research Institute, Lexington, KY) as previously described by Kenche et al. (24).

### Plasmid Construction

*mmu-mito-ncR-LDL805* (GAATTGATCAGGACATAGGGTTTGATAGTTAATATTATATGTCTTTCAAGTTCTTAGTGTTTTTGGGGTTTGGC) and *hsa-mito-ncR-LDL805* (GAACGTGTGGGCTATTTAGGCTTTATGGCCCTGAAGTAGGAACCAGATGTCGGATACAGTTCACTTTAGCTACCC) sequences were cloned into pZW1-Sno Vector (Addgene plasmid #73174; deposited by Dr. Ling-Ling Chen) between the HindIII and EcoR1 sites.

### Vector Transfection

Electroporation was carried out using a Bio-Rad Gene Pulser X cell, at 250 V and 25 μF, with 2×10^6^ cells/5 μg DNA/100 μl Gene Pulser electroporation buffer (Bio-Rad) in a 0.2-mm cuvette. Transfection efficiency was calculated as a ratio of the number of fluorescent cells to the number of Hoechst-stained cells.

### Fluorescence Confocal Microscopy

A Leica SP8-HyVolution laser scanning confocal microscope (Leica Microsystems GmbH, Wetzlar, Germany) equipped with an HCX PL APO 63×/1.4 NA and 100× oil immersion lens objectives, HyD detectors. Images were acquired at 1,256 pixels × 1,256 pixels and a step size of 0.15 μm with Leica Application Suite X software. All images were acquired using identical microscope settings and further processed using FIJI (ImageJ) and Adobe Photoshop CS5.1 software.

### Spatial Relationship Analysis

The spatial relationship of staining for *mito-ncR-LDL805*, citrate synthase, and nuclei was performed by using single-plane images with maximal signal intensities and the line scan function of LAS X software (Leica Biosystems). The plot profile tool was used to determine the fluorescence intensities of different channels along the line scan, which were plotted as a function of distance in pixels (*x* axis) and intensities (*y* axis).

### Mitochondrial Bioenergetics Analysis

Cells transfected with pZW1 vectors were seeded at 10,000 cells/well. Mitochondrial energetics was measured using a Seahorse XFe96 extracellular flux analyzer and the XF Cell Mito Stress kit (Seahorse Bioscience) according to the manufacturer’s instructions. Respiration values were calculated as described before (19).

### LP-TATs Preparation and Characterization

Nucleus targeting lipid nanoparticles containing *mmu-mito-ncR-LDL805* (LP-TAT-mmu-mito-ncR^-^LDL805) and empty lipid nanoparticles (LP-TAT) were first prepared by complexing or not *mmu-mito-ncR^-^LDL805* with cationic lipid nanoparticles consisting of DOTMA (1,2-Di-O-octadecenyl-3-trimethylammounium propane chloride salt), cholesterol and PEG (D-α-tocopherol polyethylene glycol 1000 succinate) at the DOTMA: cholesterol: PEG molar ratio of 49.5:49.5:1 through electrostatic interaction. LP-mmu-mito-ncR-LDL805 of different mass ratios of lipid:RNA were prepared as specified in figure legends. The TAT peptides (YGRKKRRQRRRC) were conjugated with Mal-PEG-DSPE [1,2-distearoyl-sn-glycero-3-phosphoethanolamine-N-[maleimide (polyethylene glycol)-2000] (ammonium salt)], and then incorporated on the surface of LP-mmu-mito-ncR^-^LDL805 at the TAT:lipid molar ratio of 1:4000 via a post-insertion process. The size, size distribution and zeta potential of TAT-LP-mmu-mito-ncR-LDL805 were measured using the Malvern ZetaSizer NanoZS (Malvern Instruments).

### LP-TATs Uptake by the Cells

Cells were replated, allowed to adhere overnight, and incubated with specified LP-TATs diluted to the final concentration of 5 nM of *mmu-mito-ncR-LDL805*, in media without FCS for 6 hours, washed and supplemented with regular media.

### Human Tissue Samples

The protocols to obtain and use de-identified human normal and COPD lung tissue specimens were approved by the Institutional Review Boards at the University at Buffalo, Case Western Reserve University, and University Hospitals of Cleveland (UB STUDY00005756, CWRU TRRC 3518 and UH STUDY20180922). Samples were obtained from The Human Tissue Procurement Facility of University Hospitals of Cleveland, embedded in paraffin blocks, and cut into 4-μm sections.

### BaseScope Duplex ISH

*hsa-mito-ncR-LDL805* was detected as previously described (19) with probes for prosurfactant C mRNA to identify AEC2s. A BaseScope duplex *in situ* hybridization (ISH) kit (ADC) was used according to the manufacturer’s instructions, with slight changes. Briefly, probes BA-Hs-LDL-1-1zz-st-C1 and BA-Hs-SFTPC-3EJ-C2 generated by the ACD probe design algorithm were used. The length of the amplification step AMPXX was increased to2 h, and that of AMPYY was decreased to 15 min. VectaMount (Vector Labs) mounting medium was used.

Levels of *hsa-mito-ncR-LDL805* expression were scored using a modified four-tier scoring system previously used for semiquantitative microscope evaluation of RNAScope: 0, no spots; 1, few spots; 2, moderate number(<10) of spots per cell; 3, high number (>10) of spots per cell (25).

### Analysis of *hsa-mito-ncR-LDL805* Staining

Images were imported to ImageJ software for analyses. Color deconvolution separated cyan (for transcripts) and magenta (for prosurfactant C) channels. Images in the cyan channel were converted to 8 bit, grayscale images, and black and white signals were inverted. The threshold of the area of interest was set as 220 (maximum, 255). The numbers and the areas of interest were counted as the number of pixels in the image that overlap with the signals for prosurfactant C in AEC2s in the magenta channel. The results were listed in a two-column table, where “area” indicated the size of the dots (pixels) and “count” indicated the number of AECs (identified as signal areas in the magenta channel). The statistics and data visualization were conducted via dplyr and ggplot2 packages in R.

### Statistical Analyses

Unless otherwise specified, data were expressed as means and standard deviations from at least three independent experiments. Student’s *t* tests were used to compare two groups, and ANOVAs with Tukey’s *post hoc* tests were used to compare multiple groups. *P* values of ≤0.05 were considered statistically significant.

## Results

### Identification and Characterization of *hsa-mito-ncR-LDL805*

The human *mito-ncR-LDL805* transcript (*hsa*-*mito*-*ncR-LDL805*) was identified from the small ncRNA library prepared from human primary AEC2s according to evolutionary conservation of its 20-nucleotide functional bit (22) (colored red in the murine sequence and yellow in the human sequence in **Figure 1A**). No nuclear mitochondrial transcript sequences corresponding to *hsa*-*mito*-*ncR-LDL805* were identified by bioinformatics analysis (**Figure 1B**) *hsa*-*mito*-*ncR-LDL805* is located at coordinates 16,472–16,546 of the D-loop of the light strand of mtDNA (**Figure 1C**). *hsa-mito-ncR-LDL805* was enriched in RNase-treated mitochondrial fractions of cell extracts, and levels were dramatically decreased in cells treated with inhibitors of mitochondrial transcription (rhodamine 6G) (**Figure 1D**). The FISH signal for *hsa-mito-ncR-LDL805* overlapped or juxtaposed the immunofluorescence signal for the mitochondrial matrix protein citrate synthase (**Figure 1E-I**; see also Figure E1 in the online data supplement). The *hsa-mito-ncR-LDL805* dots ranged from 0.2 to 1.88 μm in diameter; the larger dots likely represent several transcripts, as seen in mouse cells (19). These data indicate that the *hsa*-*mito*-*ncR-LDL805* is expressed in human primary AEC2s and is of mitochondrial origin.

**Figure 1.**
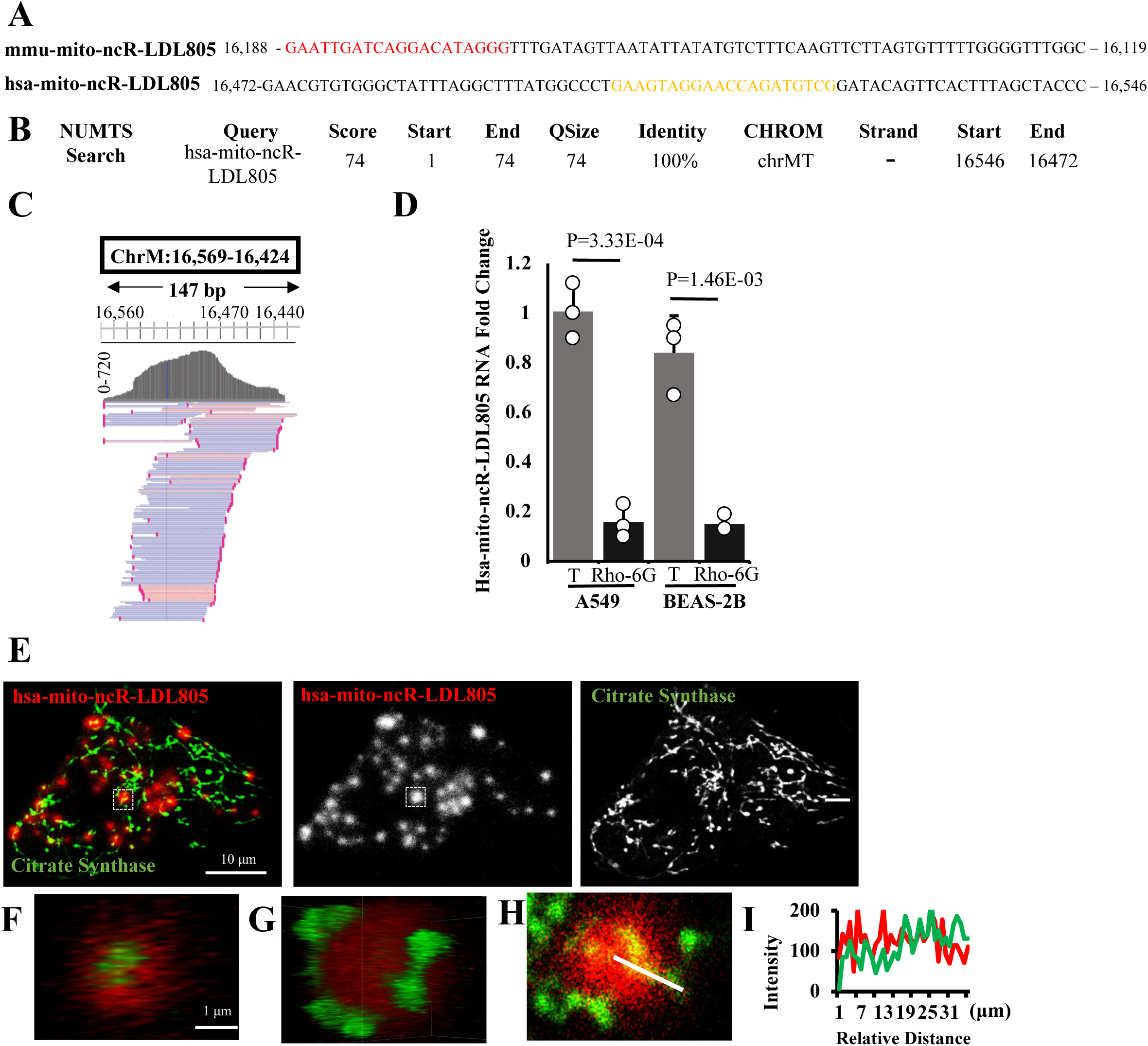
Identification of *hsa-mito-ncR-LDL805*. (**A**) Mitochondrial coordinates and sequences of murine (*mmu*) and human (*hsa*) *mito-ncR-LDL805*. The evolutionarily conserved “functional bit” is shown in red and yellow in the murine and human transcripts, respectively. (**B**) A search of nuclear mitochondrial sequences (NUMTS) conducted using BlastN at genome.ucsc with *hsa-mito-ncR-LDL805* sequence as a query, produced no nuclear hits with *hsa-mito-ncR-LDL805* as an input sequence. (**C**) Transcription alignment profile of mtDNA region 16,560–16,440. Transcripts generated by the light strand of mtDNA are represented as blue lines and those by the heavy strand are represented in red. (**D**) Expression of *hsa-mito-ncR-LDL805* evaluated by RT-qPCR from BEAS-2B and A549 cells treated or not with an inhibitor of mitochondrial transcription (rhodamine 6G [Rho-6G]) for 72 h (**E**) Confocal images of FISH for *hsa-mito-ncR-LDL805* and immunofluorescence for citrate synthase in human primary AEC2s. Specificity of the FISH probe was confirmed in samples treated with DNase (no change in signal intensity) and RNase (no signal) (see Figure E1 in the online data supplement). Magnification, ×100. The HuVolution function of the Leica SP8-HyVolution laser scanning confocal microscope was used to acquire images. Enlarged three-dimensional reconstitutions of the boxed region in panel F showing the cross-sectional distribution (**G**) and spatial localization (**F**) of citrate synthase and *hsa-mito-ncR-LDL805*. (**H**) Single two-dimensional plane of the boxed region in panel F. (**J**) Analysis of the line scan shown in panel I.

### Nuclear Translocation of *hsa-mito-ncR-LDL805* in Primary AEC2s Treated with Cigarette Smoke Extract

We assessed the levels and subcellular localization of *hsa-mito-ncR-LDL805* via FISH in primary AEC2s exposed to cigarette smoke extract. The intensity of the FISH signal increased within 2 h of exposure to cigarette smoke extract (**Figure 2A,B**). Furthermore, FISH signals were detected in the nucleus at this time point (**Figure 2A,D**). The numbers of dots and the intensity of the FISH signals decreased with longer cigarette smoke extract exposure, but this was accompanied by an increase in the nuclear presence of the signal at 6 h (**Figure 2A–F**, boxes 3 and c in panels D and E, respectively, with quantification on the right in panel F). The nuclear localization was transient and was associated with swelling of the stressed cells (**Figure 2A,C**, and **D–F**, compare boxes 3 and c [at 6 h] to boxes 4 and d [at 10 h]). The mitochondrial localization pattern was restored at 24 h when stress was resolved, as evaluated by a decrease in cell swelling (**Figure 2C**) and morphology (**Figure 2A**). Moreover, the intensity of the FISH signal at 2–10 h of exposure decreased to baseline levels 24 h after the exposure to cigarette smoke extract. These data reflect the transient response of the human transcript in human cells exposed to cigarette smoke extract.

**Figure 2.**
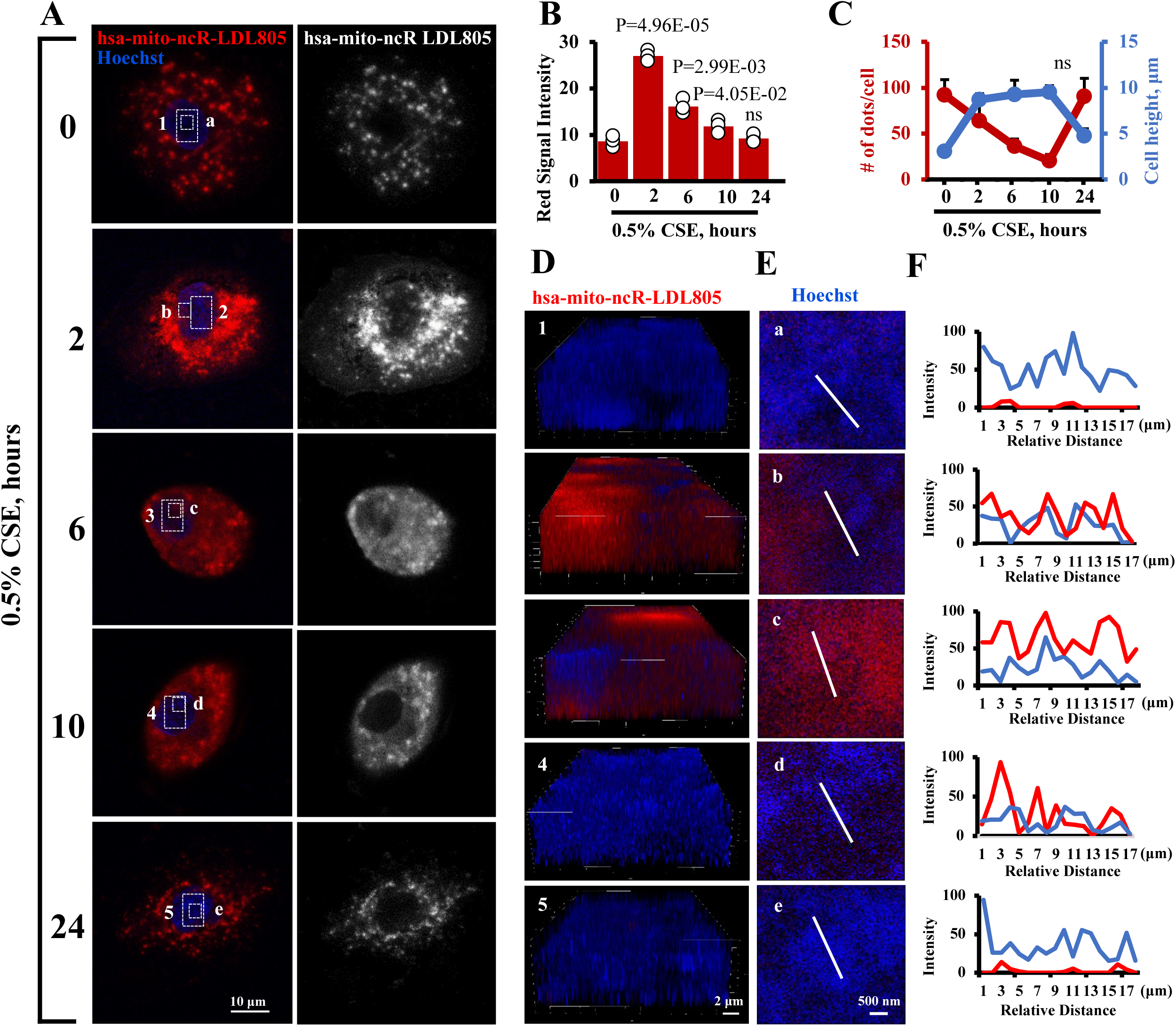
Levels and nuclear localization of *hsa-mito-ncR-LDL805* transiently increase in human primary AEC2s exposed to cigarette smoke extract. Human primary AEC2s were cultured for 24 h before being exposed to cigarette smoke extract for the indicated times. (**A**) Subcellular distribution of *hsa-mito-ncR-LDL805* was evaluated via FISH. Nuclei were visualized by Hoechst. Images represent a single plane of a maximal intensity projection of the signal for *hsa-mito-ncR-LDL805*. (**B**) Whole-cell intensity analysis of FISH signal (*n* = 3 cells per time point). (**C**) Quantification of the numbers of *hsa-mito-ncR-LDL805* dots per cell (red trace) (*n* = 12–15 cells per time point). *P* values vs 0 h: *P* = 5.34E-02 at 2 h, *P* = 2.93E-04 at 6 h, *P* = 5.26E-05 at 10 h. Quantification of cell height (blue trace) as measure of exposure-related swelling (*n* = 12–15 cells per time point). *P* values vs 0 h: *P* = 2.92E-07 at 2 h, *P* = 4.42E-10 at 6 h, *P* = 9.17E-04 at 10 h, *P* = 7.99E-04 at 24 h. Three-dimensional reconstitutions (**D**) and line scans (**E**) of boxed areas 1–5 and a–e in panel A. Confocal images were acquired with ×63 magnification and ×2.35 zoom. (**F**) Line scan analyses of areas a–e plotting the intensities of FISH (red) and Hoechst (blue) signals. For each experiment, 3–5 cells for each condition were analyzed, with at least 5 line scans per cell.

### Effects of Nuclear Delivery and Retention of *mito-ncR-LDL805*

We next sought to determine whether nuclear localization of *mito-ncR-LDL805* induces transcription of nuclear-encoded mitochondrial proteins (NeMitos) and enhances mitochondrial bioenergetics over different time periods (**Figure 3A**). Nuclear retention of *mito-ncR-LDL805* transcripts was achieved by placing the *mito-ncR-LDL805* sequence between two small nucleolar RNA genes of pWZ1-vector (26) (**Figures 3B and 4A**). The constructs were expressed in mouse (MLE12) and human (BEAS-2B) epithelial cells (**Figures 3C,D and 4B,C**, respectively). Nuclear overexpression of *mmu-mito-ncR-LDL805* in MLE12 cells and *hsa*-*mito-ncR-LDL805* in BEAS-2B cells increased transcripts levels of known NeMitos (19) (**Figures 3E and 4D**, respectively), enhanced mitochondrial bioenergetics, demonstrated as increased OCRs and higher maximal and spare respiratory capacities (**Figures 3F,G and 4E,F**), and promoted cell proliferation (**Figures 3H** and **4G**). Full-length orthologues increased mitochondrial respiration similarly to the functional bits in both cell types (see Figure E2). These data suggest that the murine and human transcripts function similarly in cells from their respective species.

**Figure 3.**
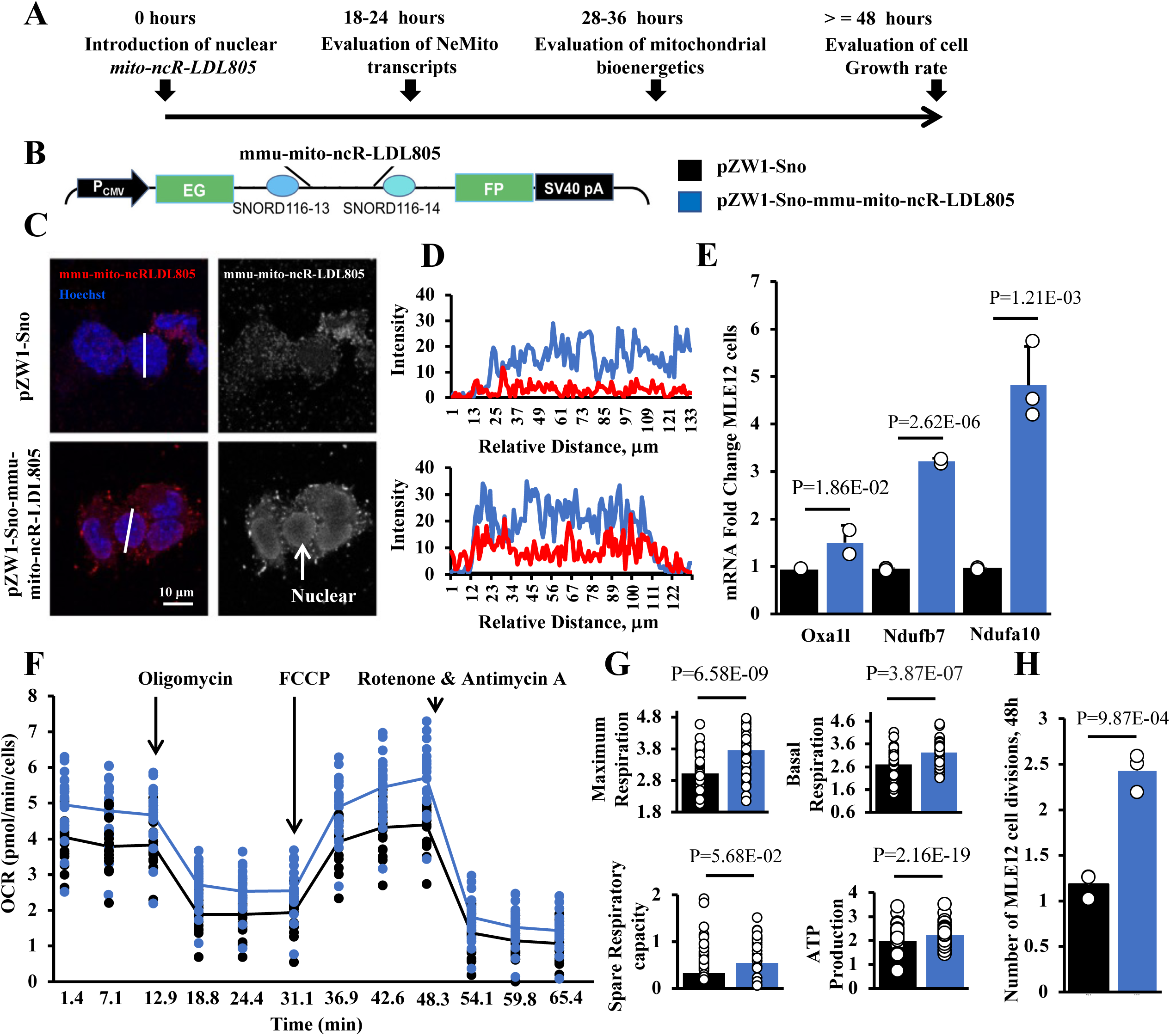
Function of nucleus-retained *mmu-mito-ncR-LDL805* in MLE12 cells. (**A**) Experimental timeline: cells were transfected with either the pZW1-Sno-*mmu-mito-ncR-LDL805* vector or the pZW1-Sno control vector. (**B**) Schematic of the pZW1-Sno-*mmu-mito-ncR-LDL805* vector. (**C**) Subcellular distribution of *mmu-mito-ncR-LDL805* 24 h post-transfection. Arrow points to *mmu-mito-ncR-LDL805* in the nucleus. Images are representatives from three independent experiments. (**D**) Analyses of the line scans indicated in panel C; red traces indicate intensity of transcript, and blue traces indicate intensity of Hoechst nuclear stain. For each experiment, 3–5 cells for each condition were analyzed, with at least 5 line scans per cell. (**E**) Expression of three representative NeMitos previously shown to be increased in cells with high levels of *mmu-mito-ncR-LDL805* : RT-qPCR analyses at 24 h post-transfection. *P* values were calculated from at least three biological experiments. **(F)** Mitochondrial bioenergetics were measured at 36 h post-transfection by using a Seahorse XFe96 extracellular flux analyzer. Oxygen consumption rate (OCR) was measured at the basal level and with subsequent sequential additions of oligomycin (1 mM), FCCP (1 mM), and rotenone (1 mM) + antimycin A (1 mM). (**G**) Quantification of bioenergetics from at least three biological repetitions of 10–12 experimental replicates each. (**H**) Cell numbers were counted 24 and 48 h post-transfection to determine the number of divisions (ratio of number of cells at 24 h to the number of cells at 48 h). Data in the graphs are representative of at least three independent experiments.

**Figure 4.**
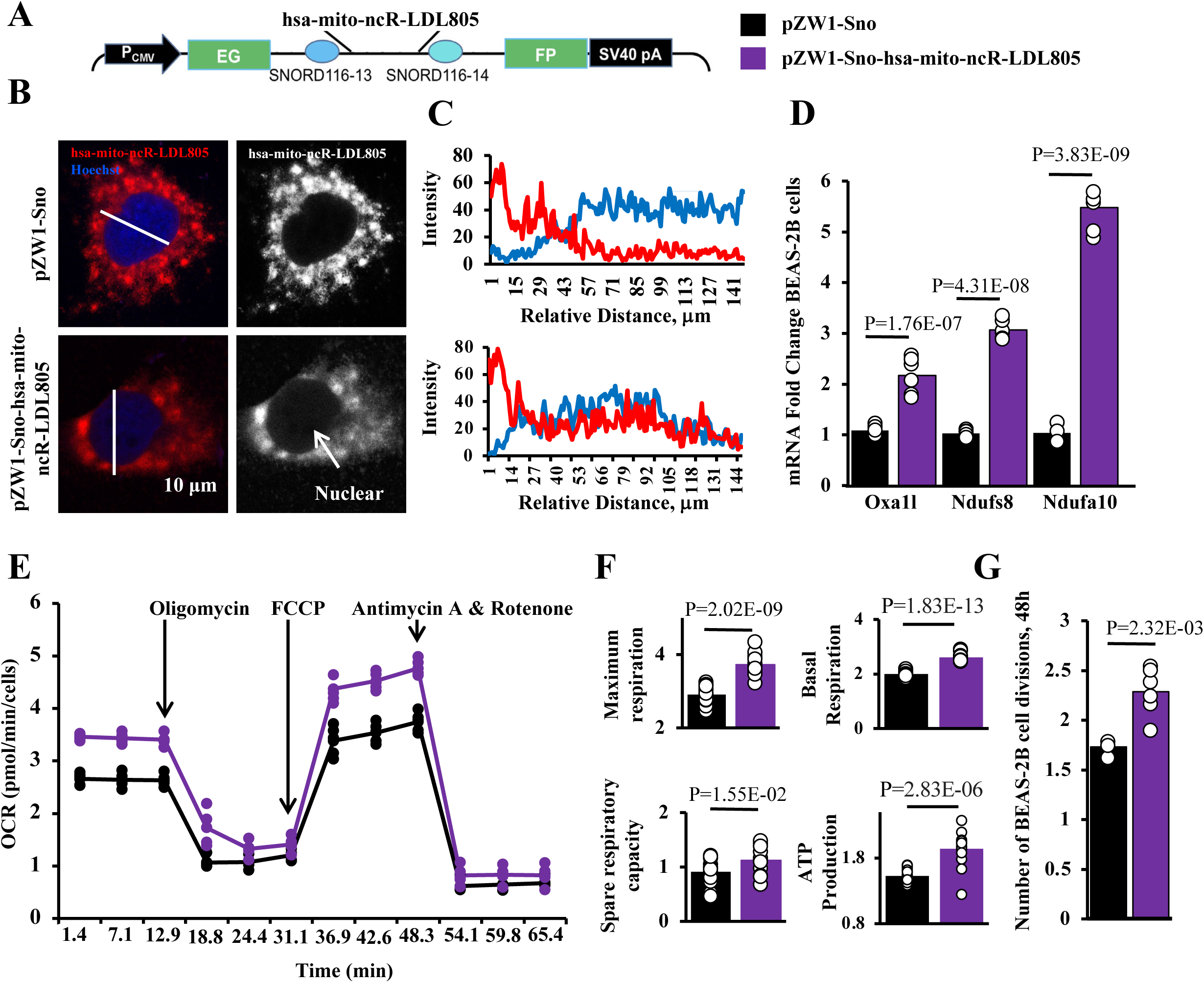
Function of nucleus-retained *hsa-mito-ncR-LDL805* in BEAS-2B cells. BEAS-2B cell were transfected with either the pZW1-Sno vector or pZW1-Sno-*hsa-mito-ncR-LDL805* vector. (**A**) Schematic of the pZW1-Sno-*hsa-mito-ncR-LDL805* vector. (**B**) Subcellular distribution of *hsa-mito-ncR-LDL805* 24 h post-transfection. Arrow points to *hsa-mito-ncR-LDL805* in the nucleus. Images are representatives from three independent experiments. (**C**) Analyses of the line scans indicated in panel B; red traces indicate intensity of transcript, and blue traces indicate intensity of Hoechst nuclear stain. (**D**) Expression of three representative NeMitos: RT-qPCR analyses at 24 h post-transfection. *P* values were calculated from at least three biological experiments. Mitochondrial bioenergetics were measured at 36 h post-transfection by using a Seahorse XFe96 extracellular flux analyzer. (**E**) Oxygen consumption rate (OCR) was measured at the basal level and with subsequent sequential additions of oligomycin (1 mM), FCCP (1 mM), and rotenone (1 mM) + antimycin A (1 mM). (**F**) Quantification of bioenergetics from at least three biological repetitions of 10–12 experimental replicates each. (**G**) Cell numbers were counted 24 and 48 h post-transfection to determine the number of divisions (ratio of number of cells at 24 h to the number of cells at 48 h). Data in graphs are representative of at least three independent experiments.

We also developed lipid nanoparticles with TAT peptides (YGRKKRRQRRRC) on the surface for targeting to the nucleus (27,28). These LP-TATs encapsulated *mito-ncR-LDL805* transcripts (**Figure 5A**; see also **Figures E3**). The small size (≤80 nm) of the particles enabling nuclear penetration was achieved by modified packaging of RNA and titration of the lipid:RNA mass ratio (compare **Figures 5B** and **E3A-C**). Nuclear localization of *mmu*-*mito-ncR-LDL805* delivered by LP-TATs in MLE12 cells was verified by FISH (**Figure 5B,C**, and **E3C**). **Figures 5** and **E3** in the online data supplement show that only LP-TATs with the lipid:RNA mass ratio of 15:1 efficiently delivered *mmu*-*mito-ncR-LDL805* into the nuclei of treated cells. Similar to SnoVector-mediated nuclear retention, LP-TAT-based nuclear delivery of *mmu-mito-ncR-LDL805* in MLE12 cells increased the levels of NeMito transcripts (**Figure 5D**), enhanced mitochondrial respiration, with increased OCRs and maximal and spare respiratory capacities (**Figure 5E,F**), and increased cell division (**Figure 5G**). These effects were observed only with LP-TATS small enough to enter the nucleus (**Figure E3**). Altogether, these data strongly support the idea that nuclear localization of *mito-ncR-LDL805* drives adaptive mitochondrial bioenergetic responses.

**Figure 5.**
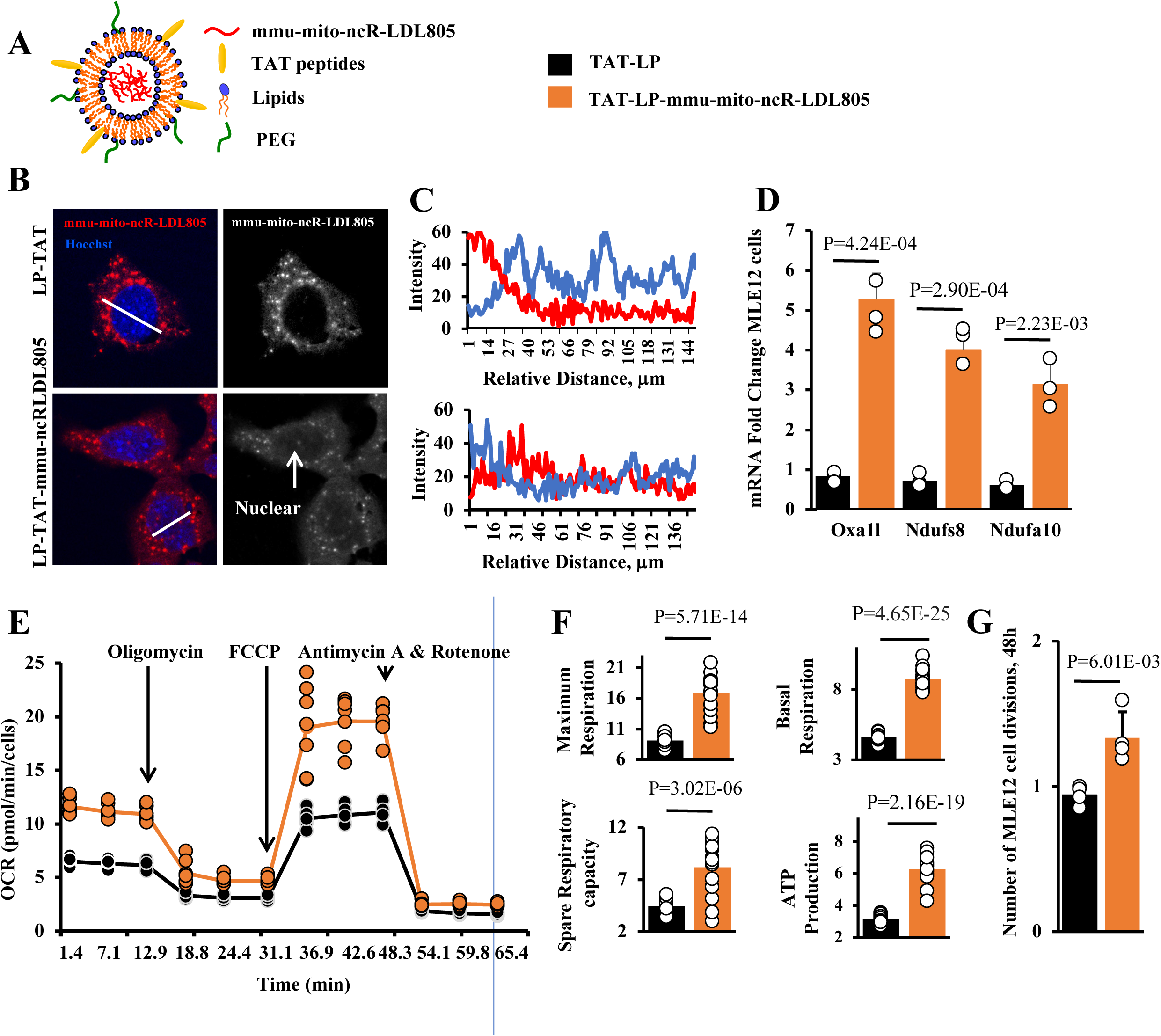
Nuclear delivery of *mmu-mito-ncR-LDL805* via LP-TATs trigger mitochondrial bioenergetics in MLE12 cells. MLE12 cells were treated with either empty LP-TAT or LP-TATs loaded with *mmu-mito-ncR-LDL805*. (**A**) Schematic of LP-TAT-*mmu-mito-ncR-LDL805*. (**B**) Subcellular distribution of *mmu-mito-ncR-LDL805* 24 h after LP-TATs were added to the medium. Arrow points to *mmu-mito-ncR-LDL805* in the nucleus. Images are representative of three independent biological experiments. (**C**) Analyses of line scans indicated in panel B (red trace, transcript; blue trace, Hoechst stain). For each experiment, 3–5 cells for each condition were analyzed, with at least 5 lines cans per cell. (**D**) Expression of three representative NeMitos 24 h after addition of LP-TATs (analyzed by RT-qPCR). *P* values were calculated from at least three representative biological experiments. (**E**) Oxygen consumption rate (OCR) was measured at the basal level and with subsequent sequential additions of oligomycin (1 mM), FCCP (1 mM), and rotenone (1 mM) + antimycin A (1 mM). (**F**) Quantification of bioenergetics from at least three biological repetitions of 10–12 experimental replicates each. (**G**) Cell numbers were counted 24 and 48 h post-transfection to determine the number of divisions (ratio of number of cells at 24 h to the number of cells at 48 h). Data in graphs are representative of at least three independent experiments.

### Low Levels of *hsa-mito-ncR-LDL805* in AEC2s from Patients with COPD

Because AEC2s in human smokers can adapt to smoke exposure (9), we hypothesized that this was in part due to adaptive changes to mitochondrial respiration driven by *mito-ncR-LDL805*. With chronic exposure, however, this adaptative response may be exhausted, leading to the manifestation of COPD (29). To test this, we obtained specimens from a tissue repository at Case Western Reserve University collected from patients with COPD (74.8 ± 9.6 years of age) and age-matched non-smoker controls (71.8 ± 6.8 years); the demographic characteristics of the patients are shown in **Table 1**.

BaseScope-duplex ISH was performed using probes specific to *hsa-mito-ncR-LDL805* and to prosurfactant C mRNA to identify AEC2s. As expected, the intensity of prosurfactant C staining was lower in specimens from COPD patients than in samples from non-smokers (**Figure 6A**); nevertheless, AEC2s were clearly detectable in all samples. AEC2s from control (non-smoker) specimens had an average of 13.33 ± 8.49 *hsa-mito-ncR-LDL805* signal dots per cell, similar to that measured in COPD samples (13.28 ± 6.61) (**Figure 6B**). The average size of the dots in specimens from non-smokers was 318.95 ± 843.87 pixels, which was significantly larger than that in COPD specimens (91.18 ± 191 pixels); notably, larger dots in the upper 25 percentile in the control samples were absent in AEC2s from patients with COPD (**Figure 6C**). A population of large spots with strong staining intensity in a control specimen are indicated by arrows in **Figure 6A**. These likely represent an accumulation of several *hsa-mito-ncR-LDL805* transcripts stored in specialized structures. Similar large-dot structures were seen in primary human cells (**Figure 1F**) and in mouse cells (19). These data suggest that the abundance or cellular storage of *hsa-mito-ncR-LDL805* in AEC2s differs between healthy and COPD subjects, strongly implying that this protective pathway is dysregulated in the diseased state.

**Figure 6.**
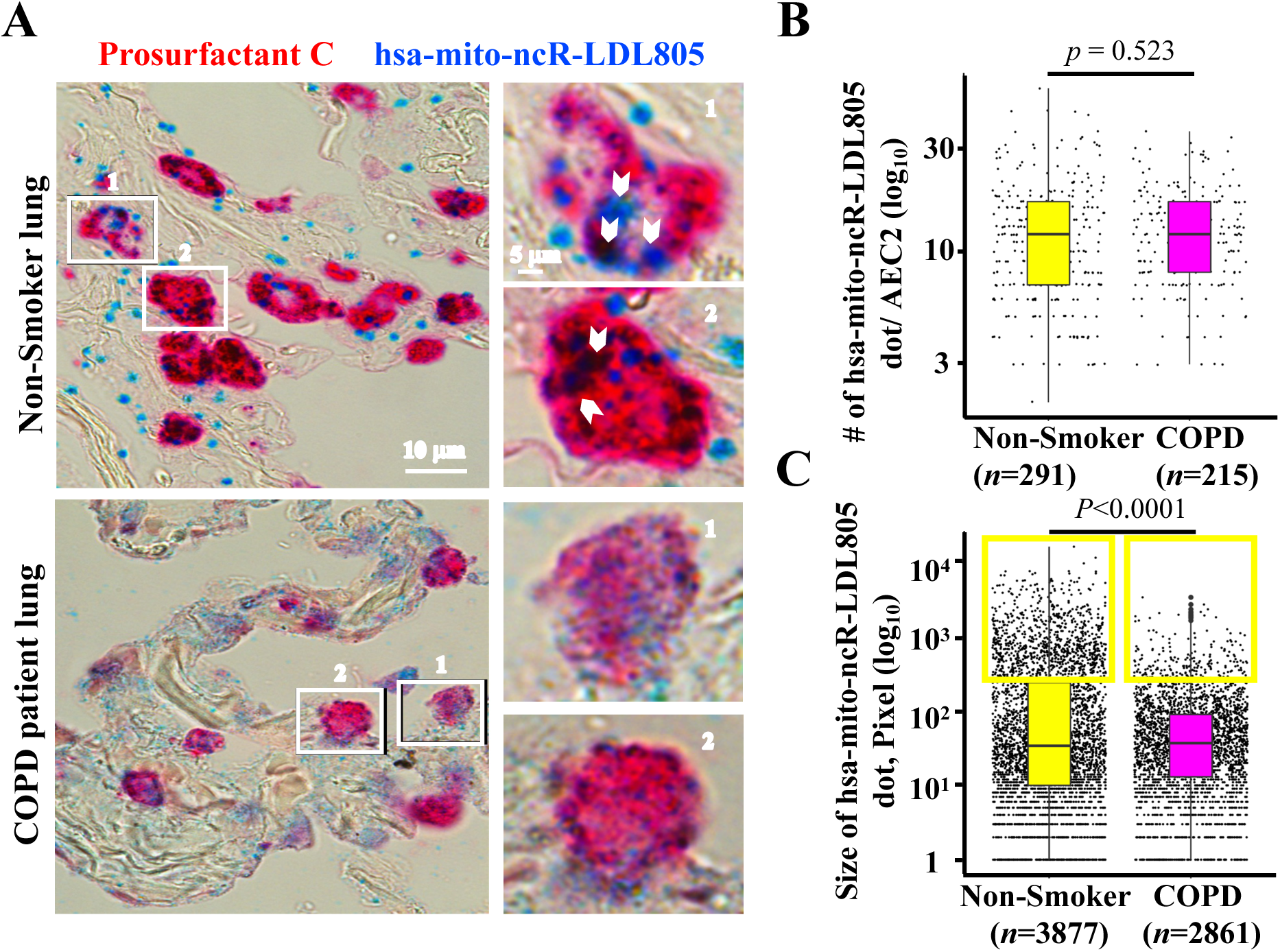
*hsa-mito-ncR-LDL805* expression is reduced in AEC2s from patients with COPD. (**A**) Specimens of human lungs obtained from non-smokers and patients with COPD were analyzed by BaseScope ISH for expression of *hsa-mito-ncR-LDL805* (blue) and prosurfactant C mRNA (red). Images were acquired using a 63× lens objective. Representative images are shown. Boxed AEC2s labeled 1 and 2 on the left are shown enlarged in panels on the right; arrows indicate large, intensely stained dots. (**B**) Quantification of the number of *hsa-mito-ncR-LDL805* spots per AEC2. (**C**) Quantification of size (pixel area) of *hsa-mito-ncR-LDL805* spots in AEC2s. For all quantifications, data were collected from five patients and five subjects; five randomly chosen areas from each specimen were analyzed: *n* = 291 AEC2s from non-smokers; *n* = 215 AEC2s from patients with COPD. The upper lines in each box plot indicate the 25 percentile of the population, the middle lines are the median or the 50th percentile of the population, and the lower lines indicate the 75th percentile. *P* values were calculated using Tukey’s *post hoc* analyses.

## Discussion

We observed a very specific mitochondrial response in AEC2s that likely underlies their resilience and proliferation in response to exposure. We provided evidence that a newly identified mitochondrial ncRNA, *hsa-mito-ncR-LDL805*, is a positive regulator of mitochondrial bioenergetics in human AEC2s via previously unrecognized mitochondria-to-nucleus signaling. The dysregulation of this response was detected in specimens from patients with COPD.

Theories of COPD pathogenesis have become more complicated as our knowledge grows, yet our ability to treat the disease remains limited. Known contributing factors include protease/antiprotease balance, chronic oxidative injury, immune responses with chronic proinflammatory signaling, and cell loss due to apoptotic, necroptotic, or other pathways of cell death (2,8–11,30). A failure of regeneration in distal airways and parenchyma, a consequence of the exhaustion of stem cell pools, was suggested recently as one of the drivers of emphysema (3). The regenerative ability of stem cells provides a level of resistance against a variety of environmental challenges, including exposure to cigarette smoke (31). However, the precise mechanism of this resilience is still being investigated. Our data suggest that adaptive mitochondrial bioenergetics are the mechanism enabling AEC2s to resist damage caused by exposure to cigarette smoke.

Mitochondria are key signaling hubs that are well integrated within the machinery for homeostasis maintenance (32). Mitochondria contribute to multiple aspects of COPD pathogenesis, such as the activation of inflammasomes, which contributes to both chronic inflammation and protease activation (10,11). Chronic mitochondrial dysfunction is accompanied by redox disbalance and bioenergetic failure, both of which accelerate cell death via energy deprivation and calcium release (33). Our findings indicate that the adaptive *mito-ncR-LDL805* response is the basis for AEC2-specific resilience and thus the absence of this results in AEC2-specific vulnerability.

Evolutionary conservation of mitochondrial ncRNAs is rare. Although the *mmu-mito-ncR-LDL805* sequence is mouse specific (according to sequence homology of the full-length transcript), a 20-nucleotide-long “functional bit” is conserved in most mammalian mitochondrial genomes (22). This conservation enabled us to identify the human homologue, which is comparable to the mouse transcript in length and location in mitochondrial genome. The studies we present here demonstrate that the conservation extends to its functions in all aspects tested so far. This suggests that the full-length transcripts likely form similar secondary structures and/or interact with similar regulatory proteins in both species and likely others, potentially revealing a novel evolutionarily conserved mitochondria-to-nucleus signaling pathway. Notably, we demonstrated via two approaches that nuclear localization of *mito-ncR-LDL805* triggers upregulation of NeMitos and increased mitochondrial respiration, which validates and extends our previous findings (22).

The way *mito-ncR-LDL805* translocates to the nucleus is not clear. However, mitochondria are known to interact with multiple intracellular organelles and structures (34), including the nucleus. Whether nuclear translocation of *mito-ncR-LDL805* requires direct interactions with those organelles is not known. Our analyses of *mito-ncR-LDL805* localization within a few hours of exposure to cigarette smoke extract suggest mitochondrial mobility and perinuclear clustering are important, possibly via the establishment of nucleus-mitochondria contact sites similar to those described for the tethering of mitochondria to the endoplasmic reticulum (35,36). Mitochondrial contacts with both the endoplasmic reticulum and the nucleus have been implicated in stress responses (32,34,35). Indeed, the outer nuclear membrane is contiguous with both rough and smooth endoplasmic reticulum [29]. Moreover, electron microscopy has shown the close association between mitochondria and the nucleus in various cell types, with stable mitochondria-associated membranes observed (37). These associations can be formed via a cholesterol-dependent mechanism that involves the mitochondrial translocator TSPO. This mechanism enables cancer cells to evade mitochondrion-induced apoptosis (37). Future studies should test if the TSPO complex tethers mitochondria to the nucleus in stressed AEC2s and if this tethering is required for nuclear localization of *mito-ncR-LDL805*.

The molecular interactions of nuclear *mito-ncR-LDL805* and its function remains to be elucidated as well. Our findings indicate that *mito-ncR-LDL805* increases the levels of NeMito transcripts. Therefore, *mito-ncR-LDL805* may function to rebuild/boost mitochondria under conditions of stress and increased energy demands. For example, the production of energy by the electron transport chain requires mitochondrion-encoded subunits, the transcription of which must match that of nuclear subunits, similar to a mechanism previously described (38). Accordingly, nuclear translocation of *mito-ncR-LDL805* was associated with an upregulation of *Oxa1l*, which encodes a mitochondrial inner membrane protein that assists in the assembly and membrane insertion of nuclear and mitochondrial subunits of the electron transport chain (39). Thus, we speculate that the bioenergetics boost provided by nuclear *mito-ncR-LDL805* generates the energy required for AEC2s to initiate cell division, “waking” them up for subsequent lung regeneration.

Our findings presented here are in agreement with results from a previous study showing that the dotted staining pattern for *mito-ncR-LDL805* disperses during stress and becomes detectable in the nucleus in mouse cells (19). Here, we further showed that this change is only transient, returning to the basal pattern after prolonged exposure to the mitochondrial stressor (e.g., cigarette smoke extract). It is possible that the reestablishment of mitochondrial *mito-ncR-LDL805* is important for AEC2 functioning, identity, and stemness under normal conditions. Our analysis of AEC2s from patients with COPD suggests that chronic smoking permanently depletes *mito-ncR-LDL805* from mitochondria and possibly nuclei. Future research will aim to clarify how dysregulation of *mito-ncR-LDL805* affects AEC2s, whether higher nuclear levels maintain healthier mitochondria, and whether mitochondrial levels are equally important.

Loss of AEC2 regeneration and exhaustion of their self-renewal capacity are factors that tip the balance towards the development of COPD. We posit that the failure to regenerate and thus exhaustion of these renewable cells is attributable to mitochondrial damage and energetic restriction. In patients with COPD and in cells with prolonged exposure to cigarette smoke extract, the mitochondrial damage likely results from the dysregulation of *mito-ncR-LDL805* in AEC2s. Therefore, we suggest that the regulation of *mito-ncR-LDL805* is a pro-survival homeostasis mechanism to maintain mitochondrial bioenergetics in AEC2s under acute stress. Further studies are needed to determine if the function of this ncRNA transcript is limited to homeostasis of AEC2s or if it also extends to other stem cells under different stressors.

Lastly, we showed that nuclear delivery of *mito-ncR-LDL805* was sufficient to induce the same mitochondrial bioenergetics responses that were triggered by exposure to cigarette smoke extract. Thus, regardless of which molecular mechanism contributes to COPD, nuclear supplementation with *mito-ncR-LDL805* is a potential strategy to enhance mitochondrial bioenergetics and delay AEC2 exhaustion. Importantly, the LP-TATs containing *mito-ncR-LDL805* in our experiments were nontoxic at the concentrations used and were optimized for both respiratory epithelium targeting and enhanced nuclear delivery (40,41). Nuclear targeted delivery was enhanced by their small size and TAT peptide modification. Small size should also protect them from internalization by alveolar macrophages (42). Therefore, such *mmu-ncR-LDL805*-containing respiratory epithelium-targeted lipid nanocarriers represent a promising strategy to improve mitochondrial function in AEC2s of patients with COPD, potentially delaying the progression of their disease.

## Acknowledgement and contributions

Special thank you to Dr. Karen Dietz for her invaluable editorial contributions, we thank Emily R. Hudson for help with developing electroporation protocol for MLE12 cells. TLM performed most of the experiments, supervised and trained MTW, analyzed data, built figures, and wrote the methods; YS performed experiments with lipid nanoparticles, analyzed the results from those studies, and generated the corresponding figures; JEB performed analysis of RNAseq data and identified human mito-ncR-LDL805, CWC analyzed the human tissues, PB helped with RT-qPCRs and quantitative analyses of *in situ* images from primary human AEC2s; MTW performed Seahorse assays with Sno-vectors; ABP, YW, MH, YP, and PAL participated in experimental design, human tissue selection, and the writing and editing of the manuscript; ABP designed the study, supervised all data analysis, and revised the manuscript with input from all authors.

This project was supported by a funding from the UBT Research Foundation to AB-P, and YP, the National Institutes of Health grants R37-DK60596 and R01-DK53307 to MH.

