## Supplementary figures and information for "Mitochondrial ncRNA LDL-805 declines in alveolar epithelial type 2 cells of chronic obstructive pulmonary disease patients"

**Supplementary data**

**Figure E1. *hsa-mito-ncR-LDL805* probe detects only RNA transcripts.** BEAS-2B cells were treated or not with either DNase, or RNase, and *hsa-mito-ncR-LDL805* specific FISH miRScope was performed. Confocal images of FISH for *hsa-mito-ncR-LDL805* Leica SP8-HyVolution laser scanning confocal microscope. All images were acquired at identical settings.

**Figure E2. Either *hsa-mito-ncR-LDL805* or *mmu-mito-ncR-LDL805* increase mitochondrial bioenergetics in both human and mouse cells.** Mitochondrial bioenergetics were measured at 36 h post-transfection by using a Seahorse XFe96 extracellular flux analyzer in (A) BEAS-2B cells and (B) MLE12 cells carrying pZW1-Sno- and either pZW1-Sno-hsa-mito-ncR-LDL805 or pZW1-Sno-mmu-mito-ncR-LDL805 vectors. Oxygen consumption rate (OCR) was measured at the basal level and with subsequent sequential additions of oligomycin (1 mM), FCCP (1 mM), and rotenone (1 mM) + antimycin A (1 mM). Quantification of bioenergetics from at least three biological repetitions of 10–12 experimental replicates each

**Figure E3. Physical characterization of different LP-TAT-mmu-mito-ncR-LDL805 and their biological effects.** LP-TAT-mmu-mito-ncR-LDL805 of different lipid: RNA molar rations were prepared and (A) their sizes, and (B) zeta potential were recorded. **(C-D)** MLE12 cells were incubated with LP-TAT-mmu-mito-ncR-LDL805 of molar ratio of 12,5:1. **(C)** Fixed and processed for FISH. Nuclear localization was not detected. Graph on the right is a representative analyses of line scan indicated in panel D (red trace, transcript; blue trace, Hoechst stain). For each experiment, 3–5 cells for each condition were analyzed, with at least 5 lines cans per cell (D) Oxygen consumption rate (OCR) was measured 36 hours post LP-TAT-mmu-mito-ncR-LDL805 addition at the basal level and with subsequent sequential additions of oligomycin (1 mM), FCCP (1 mM), and rotenone (1 mM) + antimycin A (1 mM). (**F**) Quantification of bioenergetics from at least three biological repetitions of 10–12 experimental replicates each.

**Methods:**

**RNase and DNase treatments.**

Were performed as per ACD instructions. Briefly, following the miRscope Protease digestion either RNAse A (Invitrogen Cat# AM2297) 1000 U/50 ul was added to the coverslip and incubated for 30 min at 40$^{o}$C in the HybEZ II Hybridization oven (ACD, Inc Cat# PN 321710/321720). The RNase solution was decanted, and coverslips were washed 3 times with nuclease free water. Alternatively, DNase I (Zymo Research, Cat # E1009-A) 100 units/slide of DNase in a total of 50 ul of 1X DNase I buffer (Life Technology, Catalog #: AM2222) were added to the coverslip and incubated for 30 min at 37$^{o}$C in a Cell Culture $CO_2$ incubator. The DNase solution was decanted, and cells were washed 3 times with nuclease free water. The slides were then hybridized with hsa-mito-ncR-LDL805 probe for 2 hours, at 40$^{o}$C followed by amplification steps AMP1 – AMP6 as per manufacturer instructions, except AMP5 was 15min. Red signal was developed, nuclei were visualized with Hoechst 333258 (Life Technology, cat. #P36930) (1mg/ml) and mounted with ProLong™ Gold Antifade Mounting (Life Technology, Catalog #: P36930).

**Suppl Figure E1**

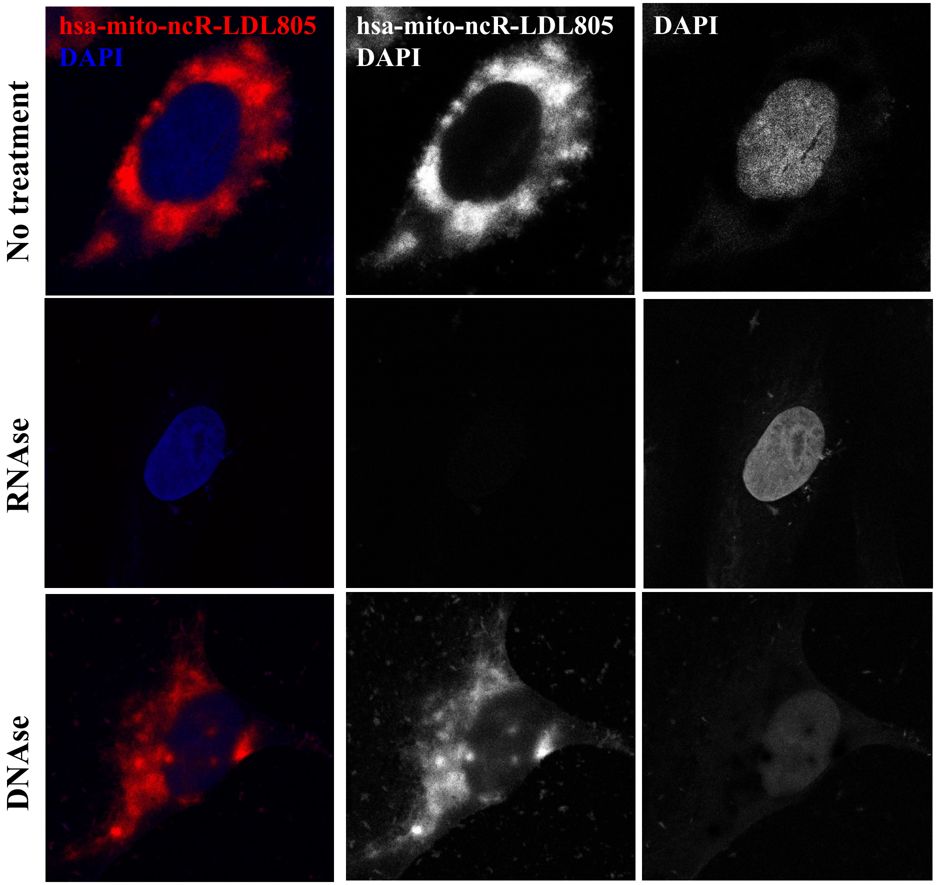

**Suppl Figure E2**

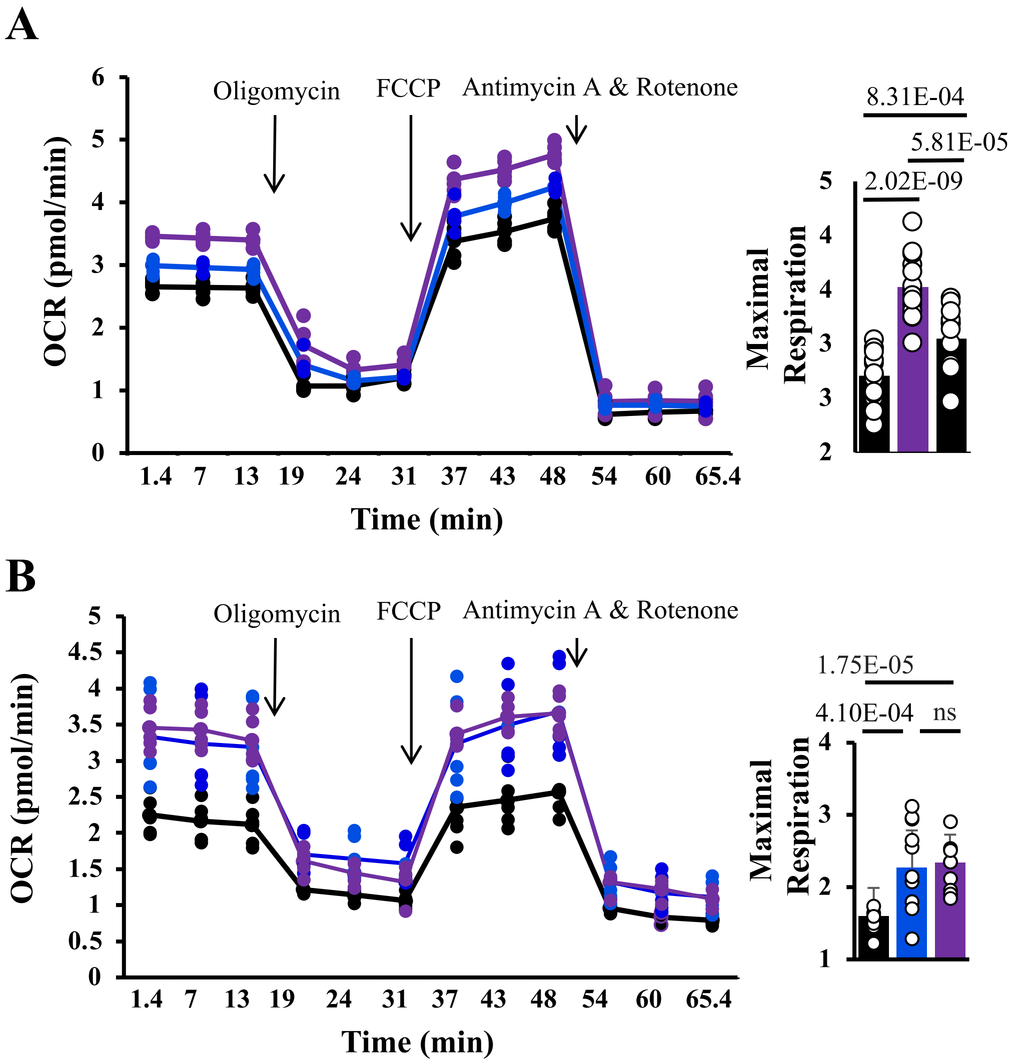

**Suppl Figure E3**

**A**

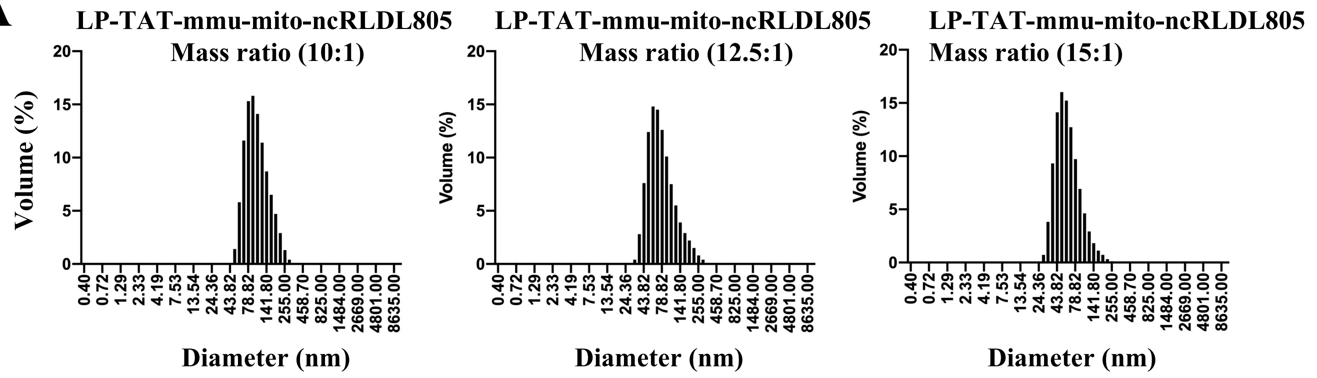


**B**

**Table 1. Size and Zeta Potential of LP-TAT-mmu-ncR-LDL805 (Average of 3 Measurements)**

| Group (Lipid:RNA mass ratio) | Volume Mean (d.nm) +/- SD | Zeta Potential (mV)+/- SD | PDI-Ave |
|---|---|---|---|
| LP-TAT-mmu-mito-ncRLDL805 (10:1) | 110.6 +/- 0.10 | 28.83 +/- 0.59 | 0.20 |
| LP-TAT-mmu-mito-ncRLDL805 (12.5:1) | 86.17 +/- 1.44 | 29.20 +/- 1.51 | 0.17 |
| LP-TAT-mmu-mito-ncRLDL805 (15:1) | 66.54 +/- -.93 | 30.17 +/- 0.25 | 0.16 |

**C** LP-TAT-mmu-mito-ncRLDL805 (12.5:1)

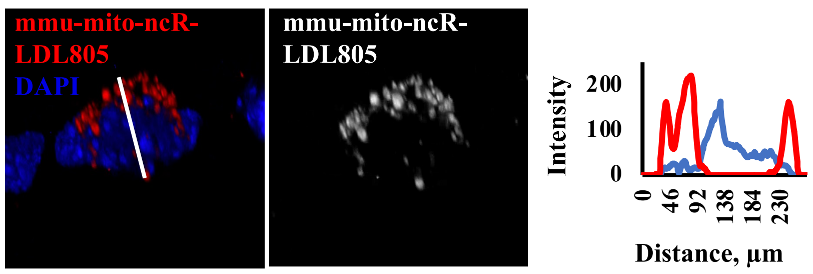


**D**

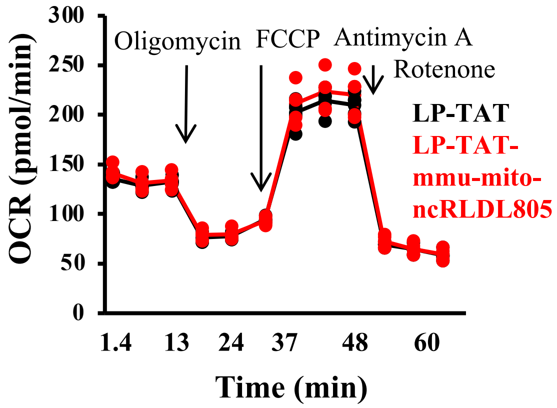
